# Multivalent, asymmetric IL-2-Fc fusions provide optimally enhanced regulatory T cell selectivity

**DOI:** 10.1101/2021.07.03.451002

**Authors:** Brian Orcutt-Jahns, Peter C. Emmel, Eli M. Snyder, Scott D. Taylor, Aaron S. Meyer

**Author notes:** **Author Emails/Contact Information:** Brian Orcutt-Jahns Peter C. Emmel Eli M. Snyder Scott D. Taylor Aaron S. Meyer.

## Abstract

The common γ-chain receptor cytokines coordinate the proliferation and function of immune cell populations. One of these cytokines, interleukin (IL)-2, has potential as a therapy in autoimmune disease but is limited in effectiveness by its modest specificity toward regulatory T cells (T_reg_s). Engineering T_reg_-selective IL-2 has primarily focused on retaining binding to the high-affinity receptor, expressed more highly on T_reg_s, while reducing binding to the lower affinity receptor with broader expression. However, other parameters, such as the orientation and valency of Fc fusion, have signaling effects that have never been systematically explored. Here, we systematically profiled the signaling responses to a panel of wild type and mutein IL-2-Fc fusions across time, cell types, and concentrations. Exploring these responses, we found that dimeric muteins have unique specificity for T_reg_s through binding avidity. A mechanistic model of receptor interactions could capture these effects and directed the design of tetravalent IL-2-Fc fusions with greater T_reg_ specificity than possible with current design strategies. Exploration of other surface targets on T_reg_s revealed that there are no other binding moieties that could be fused to IL-2 for greater selectivity. Instead, IL2Rα itself is a maximally unique surface target for T_reg_s, and so avidity is likely the only route to more selective T_reg_ interaction. However, the binding model revealed that asymmetrical, multivalent IL-2 fusions can bias avidity effects toward IL2Rα for even further enhanced T_reg_ selectivity. These findings present a comprehensive analysis of how ligand properties and their effects on surface receptor-ligand interactions translate to selective activation of immune cell populations, and consequently reveals two new routes toward therapeutic cytokines with superior T_reg_ selectivity that can be exploited for designing selective therapies in many other contexts.

**Significance Statement:** Signaling in off-target immune cells has hindered the effectiveness of IL-2 as an immunotherapy. We show that IL-2-Fc fusions with higher valency can exhibit enhanced regulatory T cell selectivity. This altered selectivity is explained by the kinetics of surface receptor-ligand binding and can be quantitatively predicted using a multivalent binding model. Using these insights, we successfully develop two new strategies for IL-2 therapies with unprecedented selectivity.

**Highlights:**

- Current IL-2 therapies are limited by a selectivity/target potency tradeoff.
- Multivalency enhances selectivity for T_reg_s through IL2Rα avidity.
- T_reg_ selectivity cannot be enhanced by targeting other surface protein markers.
- Multivalency can decouple selectivity from signaling using asymmetric cytokine fusions.

## INTRODUCTION

Cytokines that bind to the common γ-chain (γ_c_) receptor, interleukin (IL)-2, 4, 7, 9, 15, and 21, are a critical hub in modulating both innate and adaptive immune responses^1^. Each cytokine in this family binds to the common γ_c_ receptor alongside a private receptor specific for each ligand to induce signaling. These cytokines are important control mechanisms for the activity of both effector and suppressor immune populations. For example, IL-2 can increase the effector functions of CD8^+^ T cells through the upregulation of cytotoxic protein expression, as well as the suppressive capacity of T regulatory cells (T_reg_s) through increased expression of suppressive cytokines and checkpoint proteins^2–4^. Signaling through the γ_c_ family receptors also commonly results in lymphoproliferation across both suppressor and effector cell types; consequently, the γ_c_ cytokines are an important endogenous and exogenous mechanism for altering the balance of immune populations. The importance of these cytokines is observed most extremely from loss-of-function or reduced activity mutations in γ_c_, which subvert T and NK cell maturation^5^. Disruptive mutations in the private receptors can lead to more selective reductions in cell types, such as T_reg_s in the case of IL2Rα or T cells with IL-7Rα^1^. Conversely, activating mutations in these receptors, such as IL-7Rα, promote cancers such as B and T cell leukemias^6^.

Due to γ_c_ cytokines’ ability to control immune activity and abundance, they have been explored as immunotherapies in a diverse array of disease indications^7^. The most studied member of the family, IL-2, acts as both as an immunostimulant and immunosuppressant, and has been explored as a treatment for diseases ranging from cancer to autoimmunity^8– 11^. Its ability to expand T_reg_ populations, particularly at low doses, has great promise as an effective treatment for autoimmune diseases such as graft vs. host disease and hepatitis-induced vasculitis^12,13^. The efficacy of these IL-2 therapies has been hindered, however, by IL-2’s pleiotropic and non-specific activation of off-target immune populations, which simultaneously reduce therapeutic effectiveness and drive toxicities^14^. Enabling more selective activation of T_reg_s is desired to reduce these detrimental effects. However, this goal has remained elusive; effector and suppressor immune populations have only subtly-differing abundances of each IL-2 receptor subunit, and no truly T_reg_-specific marker has been discovered for targeting purposes^15^. Reducing IL-2’s affinity for IL2Rβ moderately increases T_reg_ selectivity by increasing IL-2’s reliance on IL2Rα which is found in greater abundance on T_reg_s^15,16^. However, this increase in selectivity comes at the cost of potency, as IL2Rβ is necessary for signal transduction^17^. Thus, when targeting T_reg_s, IL-2 therapies have faced a persistent tradeoff between selectivity and potency^18^.

The challenges involved in engineering simultaneously cell-selective and potent forms of IL-2 have inspired varied therapeutic designs. As mentioned, the most common approach has been to alter the receptor affinities of IL-2 to weaken its interaction with IL2Rα, IL2Rβ, or both receptors^19–23^. In most cases, the wild-type cytokine or mutein is fused to an IgG Fc to take advantage of FcRn-mediated recycling for extended half-life. Fc fusion has taken many forms such as fusion to the cytokine at the N- or C-terminus, including one or two cytokines per IgG, and including or excluding Fc effector functions^15^. So-called immunocytokines have also been employed which bind to IL-2 and block certain receptor interactions to bias signaling responses^18^. More recently, cis-targeted IL-2 fusions, such as bispecific antibodies in which a non-fused Fab binds to CD8, have been designed to deliver cytokines to cytotoxic T cells^24^. However, it is unclear whether this approach can be applied to targeting T_reg_s. These many potential therapeutic design choices necessitate design principals to direct therapeutic engineering.

Here, we systematically profile the signaling specificity effects of engineered cytokine alterations, including affinity-altering mutations and Fc-fusion formats, to map the current landscape of cell-selective IL-2 designs. Through this systematic evaluation, we identify Fc fusion valency to be an important factor in cell type selectivity. The signaling specificity of all muteins and Fc formats quantitatively matches a multivalent binding model, both between cell types and across cell-to-cell variation within a cell type, indicating that the effect of cytokine multivalency is derived from altered surface receptor binding avidity. We then use this model to demonstrate that cytokines engineered in higher valency formats are predicted to confer greater specificity towards a variety of immune cell types. These insights were experimentally validated by designing and testing two novel tetravalent IL-2 muteins in both symmetric and asymmetric forms which displayed superior T_reg_ signaling selectivity. The superior performance of asymmetric tetravalent IL-2 fusion proteins also demonstrates how bitargeting/asymmetry can decouple targeting from signaling, enabling new therapeutic opportunities. In total, our analysis and experimental findings demonstrate that cytokine valency is an unexplored direction for further enhancing selective signaling responses and that many opportunities for using multivalency engineering exist within the γ_c_ cytokine family and beyond.

## RESULTS

### Systematic IL-2 variant profiling reveals multiple determinants of response

To explore how IL-2 mutations affect signaling across immune populations, we stimulated peripheral blood mononuclear cells (PBMCs), collected from a single donor, with 13 IL-2 muteins (Fig. 1a/b, Table S1). Our panel principally included IL-2 mutants previously developed to confer enhanced-T_reg_ selective signaling^15,22,23^. In addition to changes in receptor affinities, we included variation in two structural features: Fc fusion at either the C- or N-terminus, which has been shown to alter receptor-interaction kinetics^15^, and fusion in both monomeric and dimeric formats. Stimulated cells were collected at four time points using 12 treatment concentrations. The PBMCs were then stained for canonical cell type markers and phosphorylated STAT5, a commonly used read-out of IL-2 signaling response, allowing us to separate signaling response by cell type. Four different cell types (T_reg_, T_helper_, CD8+, NK) were gated and quantified (Fig. S1a–d). T_reg_ and T_helper_ cells were further dissected into low, average, and high IL2Rα abundance by isolating subpopulations using three logarithmically spaced bins (Fig. S1j). For a surface-level visualization of the effects of time, cell type, receptor abundance, ligand format, ligand affinity, and concentration, we organized our signaling data into a heatmap (Fig. 1c). The complexity of the data demanded closer examination.

**Fig. 1.**
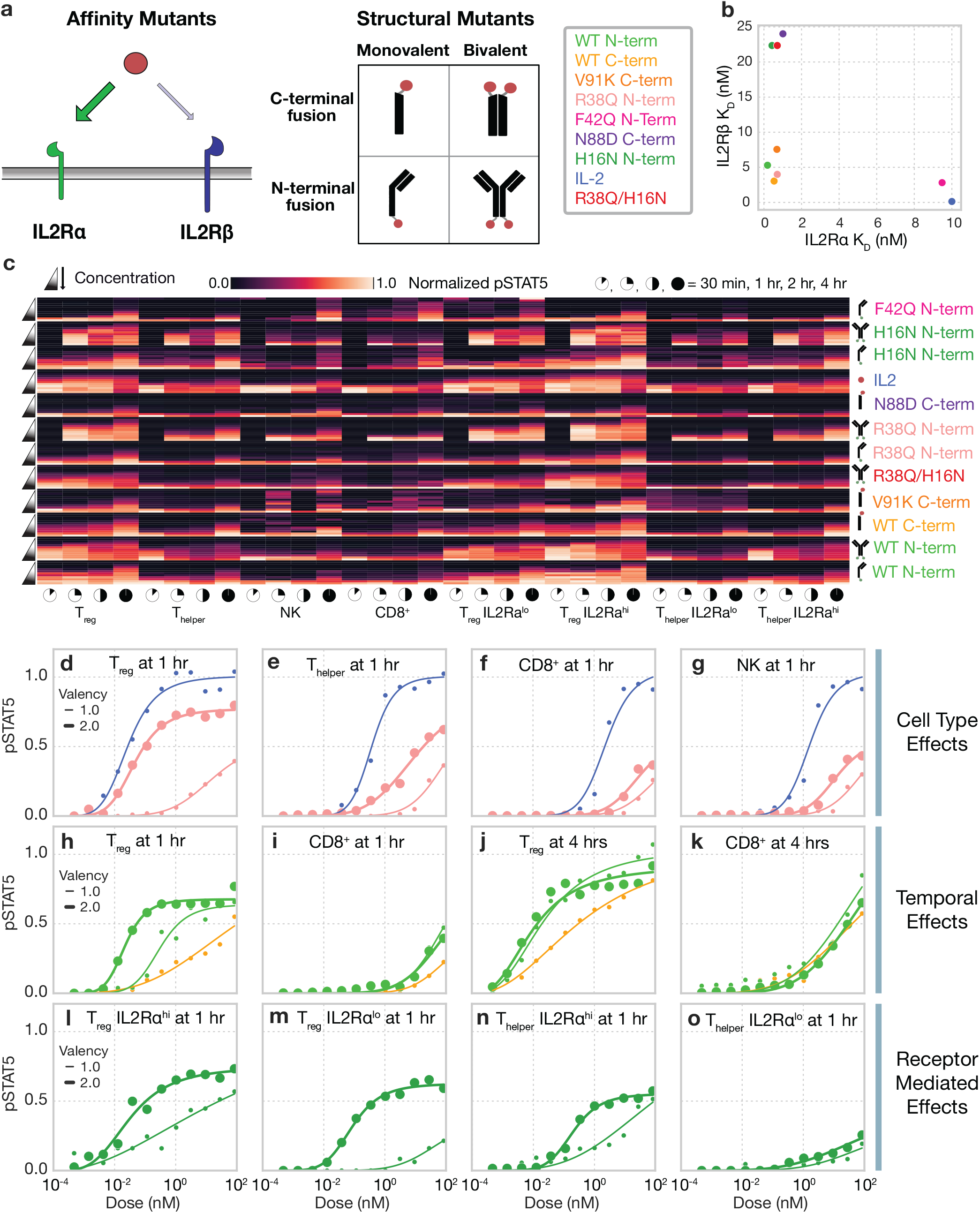
Systematically profiling IL-2 muteins reveals determinants of response. **(a)** Schematic of affinity and structural mutants explored. **(b)** IL2Rα and IL2Rβ affinities of each IL-2 variant. **(c)** Heatmap of phosphorylated STAT5 measurements for each cell type, time point, ligand, and concentration. pSTAT5 measurements are normalized for each cell type. **(d–o)** STAT5 phosphorylation response curves for immune cells stimulated with select IL-2 muteins. Time points and cell types are indicated in subplot titles.

We selectively highlighted several dose-response curves to demonstrate the importance of our comprehensive characterization (Fig. 1d–o). First, as expected, we found that the affinity with which each IL-2 interacted with receptors divided responses (Fig. 1d–g); for example, wild-type (WT) IL-2 most potently activated all cell types, as expected given it bound to IL2Rβ with the greatest affinity (Fig. 1b, d–g). Valency played also had a prominent effect on signaling response; the bivalent Fc fusion form increased sensitivity and potency of response across all cell types.

We also found that temporal dynamics affected response characteristics (Fig. 1h–k). For example, we found that C- or N-terminus Fc-fused IL-2 demonstrated distinct responses in T_reg_s at 1 hour of treatment but shared responses after 4 hours of treatment (Fig. 1h–k). Temporal effects are likely influenced by receptor mediated endocytosis of IL-2 receptor subunits and transcriptional changes enacted by IL-2 signaling and STAT5 phosphorylation^25,26^.

Finally, we found that receptor abundance interacted with cell identity to modulate response (Fig. 1l–o). T_reg_ populations with high amounts of IL2Rα strongly responded to monovalent H16N N-term, and the bivalent form moderately enhanced this response. However, in the IL2Rα^lo^ T_reg_ cells, the effect of bivalency was even greater—only bivalent H16N induced a significant response. IL2Rα^high^ T_helper_ cells also showed a moderate increase in potency with bivalency, like the IL2Rα^high^ T_reg_ cells, but the IL2Rα^lo^ population showed no distinction between the monovalent and bivalent fusions. Thus, immune populations are further subdivided by receptor abundance into subpopulations with distinct cellular responses.

In total, the dynamics of response, cell type, concentration, ligand affinity, Fc fusion valency, and Fc fusion orientation all play roles in determining cellular response. These determinants interact in unique and often unintuitive manners, requiring a more comprehensive accounting of their effects.

### Ligand valency and affinity interact to form unique cell type selectivity profiles

Given the coordinated importance of time, ligand valency, ligand affinity, cell type, and receptor expression, we next sought to focus on how ligand format affected T_reg_ selectivity. The selectivity of IL-2 for specific cell types corresponds closely to its therapeutic potency and potential toxicities^19,21,22,27^.

To better understand the relationship between T_reg_ selectivity and ligand properties, we plotted the ratio of STAT5 phosphorylation (fit by a Hill curve) in T_reg_s to that of off-target cells for each ligand across our concentration range (Fig. 2a/b/e/h). We saw that shape of each selectivity curve varied widely for each ligand and off-target cell type considered (Fig. 2b/d/f). T_reg_ selectivity quantified against CD8^+^ and NK cells most prominently separated bivalent from monovalent ligands. Since IL2Rα affinity widely varied between our IL-2 mutants and is a known regulator of T_reg_ selectivity, we sought to understand how affinity differences contribute to T_reg_ selectivity. We plotted ligand IL2Rα affinity against the peak T_reg_ selectivity observed across concentrations (Fig. 2c/e/g). Due to the high abundance of IL2Rα displayed by T_reg_s (Fig. S1i), we expected to see a positive correlation between IL2Rα affinity and peak T_reg_ selectivity. However, we saw that this relationship varied in a cell type- and valency-dependent manner. When considering either CD8^+^ or NK cells, decreasing IL2Rα affinity led to decreases in the peak T_reg_ selectivity of monovalent muteins, but also, somewhat surprisingly, little relationship with the bivalent selectivity peaks (Fig. 2c/e). When considering T_helper_ populations, which have greater amounts of IL2Rα than CD8^+^ and NK cells, we observed that decreases in IL2Rα affinity led to increases in maximum selectivity for both monovalent and bivalent muteins (Fig. 2g).

**Fig. 2.**
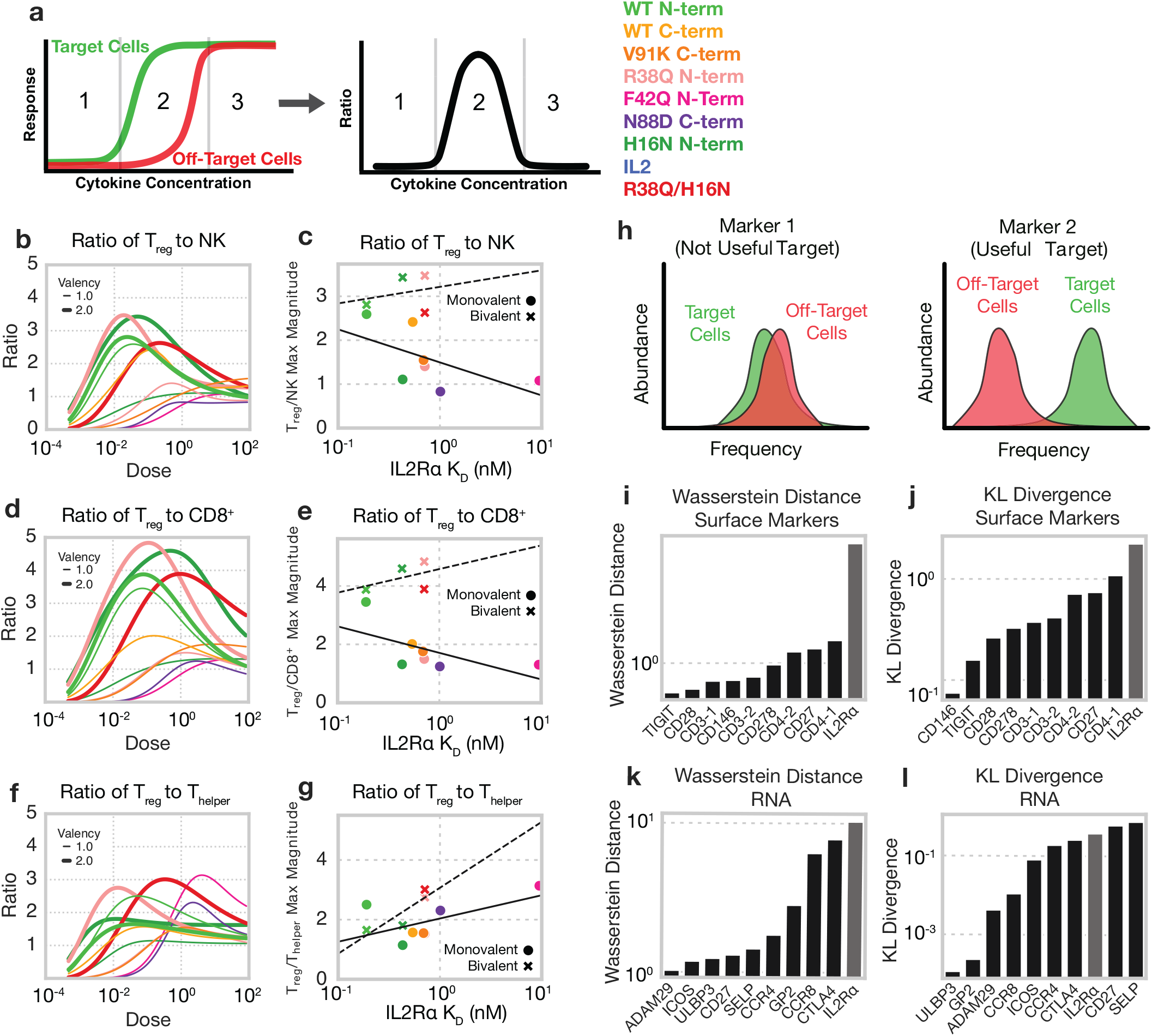
IL-2 muteins display structural- and affinity-dependent T_reg_ selectivity that cannot be overcome with cis-targeting strategies. **(a)** Schematic describing ratio of activation between target and off-target immune populations. **(b, d, f)** Ratios of T_reg_ pSTAT5 to NK (b), CD8^+^ (d), and T_helper_ (f) pSTAT5 dose response curves at 4 hours. Ratio was defined as the ratio of Hill curves fit to experimental data for target and off-target populations, as in Fig. 1. **(c, e, g)** Maximum ratio of T_reg_ to off-target population signaling vs. IL2Rα affinity. Lines of best fit were fit to monovalent (solid) and bivalent (dashed). (**h)** Schematic depicting how useful markers for conferring selectivity are selected. **(l–o)** Most unique markers for T_reg_s in CITE-seq dataset identified using Wasserstein distance (i,k), and Kullback-Leibler Divergence metrics (j,l) using surface markers (i,j) and RNA data (k,l).

Unsurprisingly, the mutein concentration at which the maximum T_reg_ selectivity occurred was higher for ligands with weaker IL2Rα affinity across all cell types (Fig. S2). In total, we show here that affinity and valency affect the selectivity profiles across ligand doses in distinct yet intertwined manners; to understand these relationships, a method mapping each of these factors simultaneously is needed.

### The protein expression and abundance profile of T_reg_s reveals limits to enhancing IL-2 T_reg_ selectivity through cis targeting

While IL2Rα is more abundant in T_reg_s, the difference is subtle compared to some off-target cells, making selectively targeted activation more challenging^15,18^. Consequently, we wondered whether a cis targeting strategy, in which IL-2 is fused to a domain binding some other T_reg_-specific surface marker, would provide even greater selectivity. To explore this possibility, we used a CITE-seq data set in which >211,000 human PBMCs were simultaneously analyzed for 228 surface markers coupled with single-cell RNA-seq^28^. Our previous work shows that specificity is conferred by markers expressed at a high ratio between target and off-target cells^29^. As a measure of difference, we calculated the Wasserstein distance and Kullback-Leibler divergence of each surface marker abundance and expression between T_reg_ populations and all off-target PBMCs. These complimentary distance metrics were chosen to reflect two different measures of difference: the Wasserstein distance is maximized when transforming one distribution to another would require changing the cells to a large degree, while the Kullback-Leibler distance is maximized when the overlap between two distributions is minimized. We were surprised to find that IL2Rα was the most consistently and uniquely expressed marker on T_reg_s by both proteomic and transcriptomic analysis (Fig 2l-o). These results were reinforced by using both a linear and non-linear classifier to identify which surface markers and transcripts were most informative for T_reg_ classification; this analysis again found that IL2Rα was optimal (Fig S3). In total, these analyses show that binding alternative surface markers cannot improve IL-2 selectivity for T_reg_s.

### Bivalent Fc-cytokine fusions have distinct cell specificity but shared dynamics

Understanding that selectivity for T_reg_s must be derived through engineering binding to its cognate receptors, we sought to develop a more complete view of the various structural choices for IL-2 fusion design. Exploring variation in response across cell types and ligand treatments is challenging due to its multidimensional nature. Restricting ones’ view to a single time point, cell type, or ligand concentration provides only a slice of the picture (Figs. 1 & 2)^15,30^. Dimensionality reduction is a generally effective tool for exploring multidimensional data. However, flattening our signaling data to two dimensions and using principal components analysis failed to help isolate the effects of concentration, ligand properties, time, and cell type (Fig. 3a). Therefore, to better resolve our data, we organized our profiling experiments into a four-dimensional tensor organized according to the ligand used, concentration, treatment duration, and cell type in the profiling. We then factored this data using non-negative canonical polyadic (CP) decomposition, a technique that represents n-dimensional tensors as additively separable patterns, themselves approximated by the outer product of dimension-specific vectors^31^. We used CP decomposition to derive factors summarizing the influence of each dimension (Fig. 3b).

**Fig. 3.**
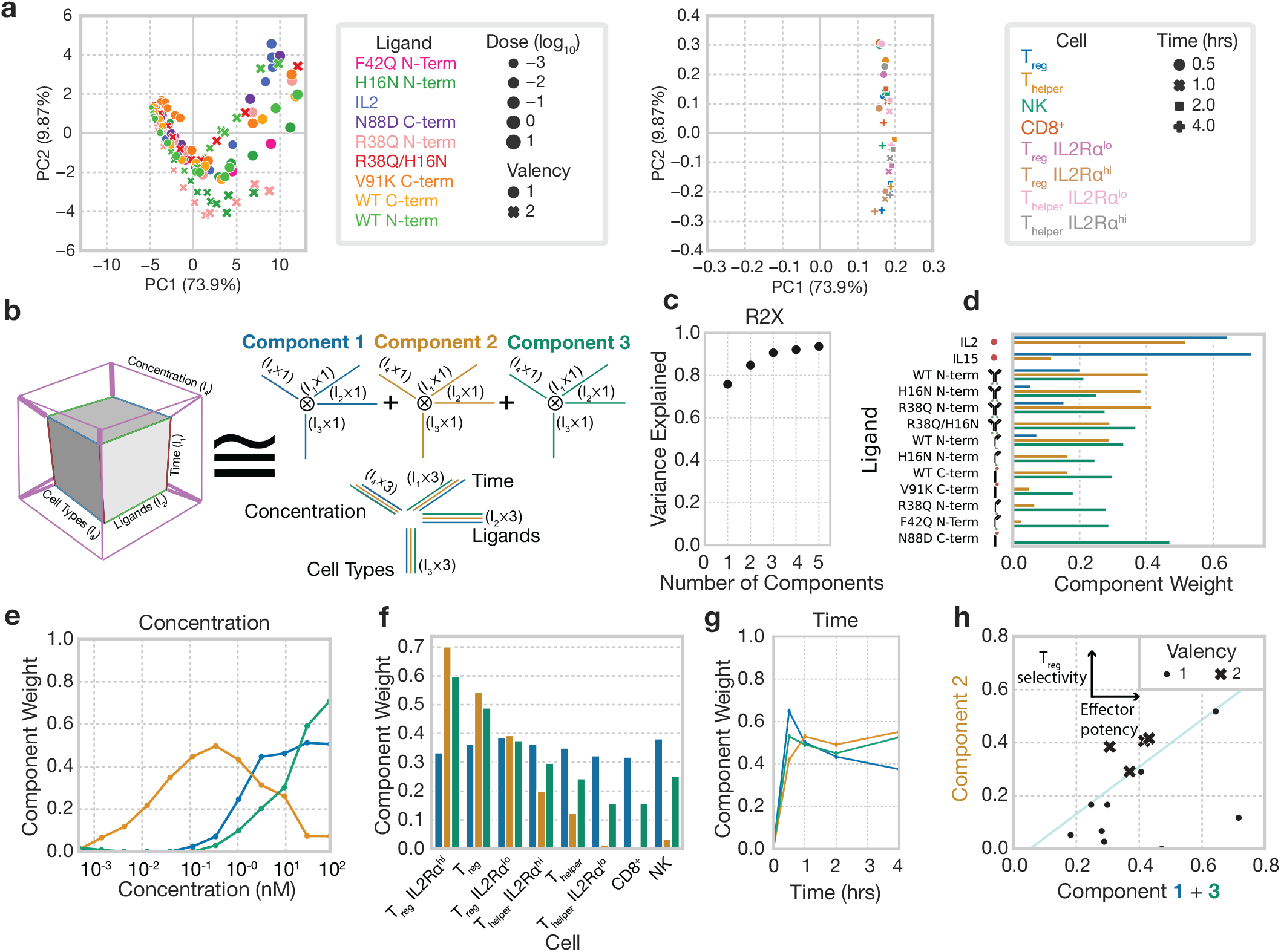
Tensor-based decomposition reveals unique selectivity defined by fusion valency. **(a)** Scores and loadings plot of pSTAT5 signaling data as calculated using principal components analysis. **(b)** Schematic representation of non-negative canonical polyadic (CP) decomposition. Experimental pSTAT5 measurements are arranged in a tensor according to the duration of treatment, ligand used, cytokine concentration, and cell type. CP decomposition then helps to visualize this space. **(c)** Percent variance reconstructed (R2X) versus the number of components used. **(d)** Component values for each IL-2 mutant. **(e)** Component values representing the effect of IL-2 concentration. **(f)** Component values representing cell type specificity. **(g)** Component values for the effect of treatment duration. **(h)** Sum of Component 1 and 3 weights (off-target cellular signaling) vs. Component 2 (T_reg_ signaling) weight for each monovalent and bivalent ligand. Thin line is included for visualization purposes only.

Three components explained roughly 90% of the variance within the dataset (Fig. 3c). Factorization separated distinct response profiles into separate components, and the effect of each dimension (e.g., time, concentration) into separate factors. For instance, component 1 almost exclusively represented responses to wild-type cytokines (Fig. 3d), which were the only ligands which were not Fc-fused, showing a response primarily at high concentrations (Fig. 3e), with broad specificity (Fig. 3f) and a signaling profile that peaking 30 minutes and then more rapidly decreasing (Fig. 3g). An alternative way to interpret the factorization results is to compare profiles within a single factor. For example, component 1 led to a less sustained profile of signaling response as compared to the other signaling patterns (Fig. 3g).

Remarkably, components 2 and 3 cleanly separated ligands conjugated in bivalent or monovalent forms, respectively (Fig. 3d/h). In fact, ligand valency was represented more prominently than differences in receptor affinity between muteins. Component 2 had uniquely high T_reg_ specificity (Fig. 3f) and was most represented at intermediate concentrations (Fig. 3e). Component 2 was also highly correlated with IL2Rα abundance in subsets of T_reg_ and T_helper_ cells, suggesting that bivalent molecule’s specificity for T_reg_s is mediated by their higher abundance of IL2Rα. Component 3 had a broader cell response (Fig. 3f) and increased monotonically with concentration (Fig. 3e). Despite these strong differences in specificities, both components had nearly identical time dynamics (Fig. 3g). While other ligand variation influenced the potency and selectivity of each ligand, only the bivalent Fc fusions, regardless of their receptor affinities, more highly weighted component 2, where T_reg_ response was highly weighted, over components 1 and 3, which represented effector cell response (Fig. 3h). In total, these results indicated that mono- and multivalent cytokines shared identical dynamics and that, while Fc fusion and affinity modulation affect response, ligand valency was a critical and prominent determinant of specificity.

### Variation in IL-2 responses is explained by a simple multivalent binding model

Having observed that T_reg_ selectivity is prominently enhanced by multivalency, we sought to determine whether cell surface binding on its own could explain these selectivity differences. To do so, we applied a two-step, equilibrium, multivalent binding model to predict IL-2 response, assuming that signaling response was proportional to the amount of active receptor-ligand complexes^32^. Within the model, ligand binding first occurs with kinetics equivalent to the single binding site, and then subsequent interactions occur proportionally to affinity, adjusted by 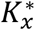, a cross-lining constant that corrects for differences between monovalent and multivalent interactions. We fit this model to our signaling experiments and evaluated its concordance with the data. The model is very simple, with the cross-linking parameter being the only non-scaling fit parameter; this parameter had an optimum at 1.2 × 10^−11^ #/cell, consistent with that seen for other receptor families^33–35^.Overall, we observed remarkable consistency between predicted and observed responses (R^2^ = 0.85), and accuracy was maintained when examining data subsets including individual cell types and ligands (Fig. 4b–c).

**Fig. 4.**
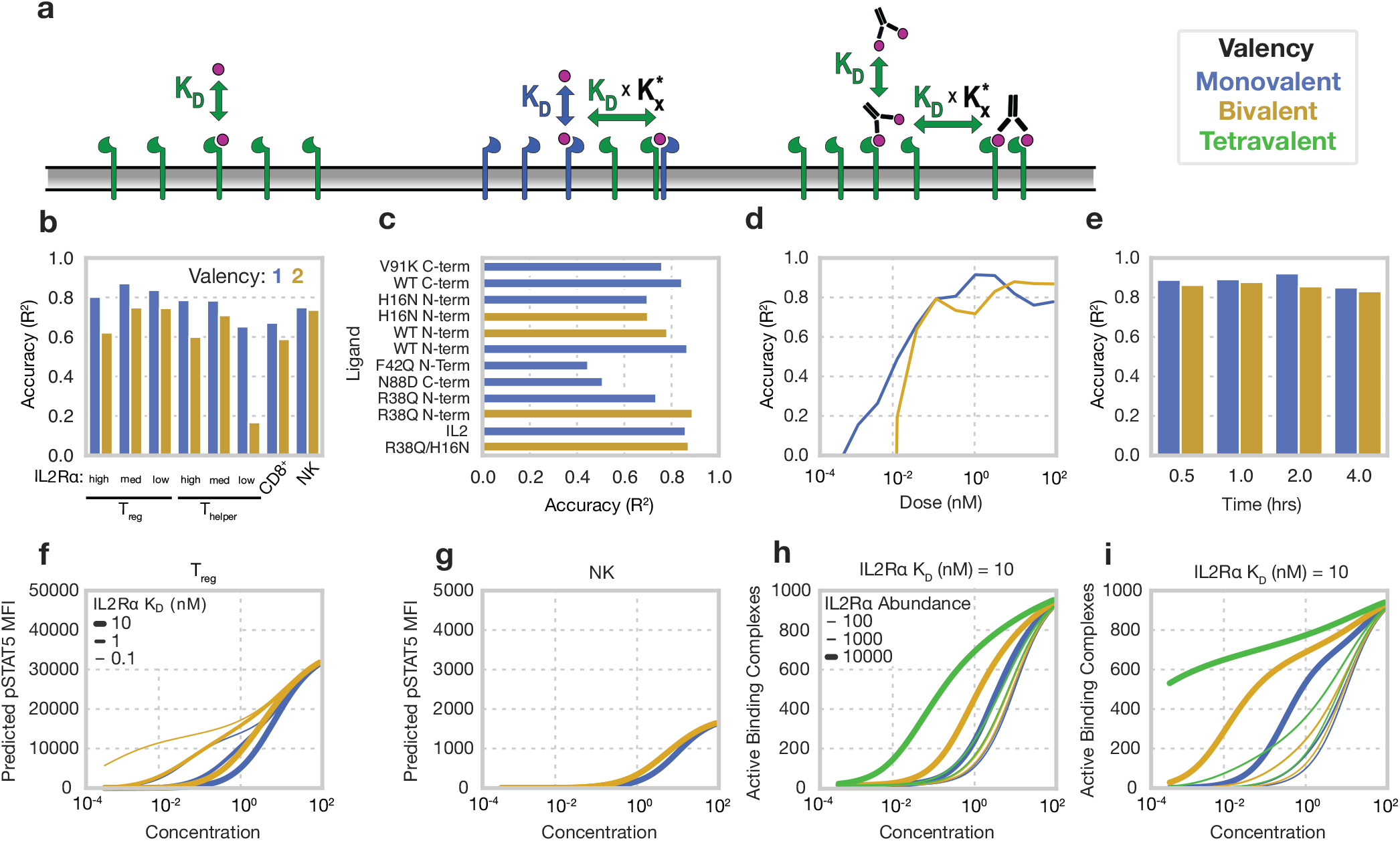
Responses are predicted by a simple multivalent binding model. **(a)** A simplified cartoon of the model. Initial association of multivalent ligands proceeds according to monovalent affinity, and subsequent binding events proceed with that affinity scaled by the K^∗^ parameter. **(b, c)** Model’s accuracy subset by cell type **(b)** and ligand **(c)** for all mono- and bivalent IL-2 muteins. **(d, e)** Model’s accuracy subset by concentration **(d)** for all ligands and time **(e)** for all ligands, concentrations, and cell types. All accuracies are calculated as a Pearson’s correlation R^2^ score for experimental cytokine responses at 30 mins and one hr. **(f, g)** Model-predicted pSTAT for T_reg_s (f) and NK cells (g) in response to mono- and bivalent IL-2 ligands with 10 nM IL2Rβ K_D_. **(h, i)** Predicted number of active signaling complexes formed on cells with 1000 IL2Rβ receptors and varying numbers of IL2Rα for ligands with affinities of 10 nM K_D_ for IL2Rβ and either 1 nM (f) or 10 nM (i) K_D_ for IL2Rα.

To ensure that our model was not simply capturing a trend towards higher signaling with increasing concentration, we examined our model’s accuracy within specific cytokine concentrations (Fig. 4d). Our model did not predict response at the lowest concentrations as there was little to no response in the data itself but increased in accuracy at concentrations where responses were observed. Finally, we examined how the model’s accuracy varied within each timepoint; initial responses (30 mins) were most accurately predicted, with a slight decrease in accuracy over longer timescales (Fig. 4e). This is expected, given that longer treatments likely involve various compensatory mechanisms such as the degradation or increased transcription of IL-2 receptor subunits^26,36^. In total, multivalent cell surface binding showed quantitative agreement with the pattern of cell-type-specific responses to IL-2 muteins, supporting that the specificity enhancement of bivalency is derived from receptor avidity effects and is explained by a simple model of cell surface binding.

Upon finding that our model was broadly predictive of cell-type specific signaling responses, we sought to use our model to understand and visualize how valency and affinity interact to determine T_reg_ selectivity. Here, our model showed that T_reg_ response is strongly governed by IL2Rα affinities and that these effects have an exceptionally strong relationship with valency, particularly at intermediate cytokine doses, while NK signaling barely varied across ligands of varying affinities (Fig. 4f/g). We then used the model to explore how receptor abundance affects multivalent ligand binding (Fig. 4h/i). Here, theoretical cell populations expressing 10^4^ IL2Rα and 10^3^ IL2Rβ molecules varied widely in their response to multivalent IL-2 muteins (Fig. 4h), while cells expressing very few IL2Rα receptors and the same abundance of IL2Rβ barely altered in their response (Fig. 4i).

These results demonstrate how multivalent cytokines with high IL2Rα affinities uniquely and selectively target T_reg_s through IL2Rα-mediated avidity effects.

### Multivalency provides a general strategy for enhanced signaling selectivity and guides the development of superior IL-2 muteins

Given that a simple binding model accurately predicted cell type-specific responses to IL-2 and that bivalent, Fc-fused IL-2 muteins have favorable specificity properties, we computationally explored to what extent multivalency might be a generally useful strategy. While monovalent ligand binding scales linearly with receptor abundance, multivalent ligands bind nonlinearly depending upon receptor abundance^37^. Thus, multivalent ligands should be able to selectively target cells with uniquely high expression of certain γ_c_ family receptors.

Valency enhancements are only apparent with coordinated changes in receptor-ligand binding affinities^29^. Therefore, we optimized the receptor affinities of simulated ligands while varying valency. We first designed IL-2 muteins of varying valency to obtain optimal T_reg_ specificity (Fig. 5a). As expected, ligand valency increased achievable selectivity past that possible using a monovalent cytokine format at any receptor affinities. Muteins of higher valency required reduced IL2Rα affinity to achieve optimal T_reg_ selectivity (Fig. 5b).

**Fig. 5.**
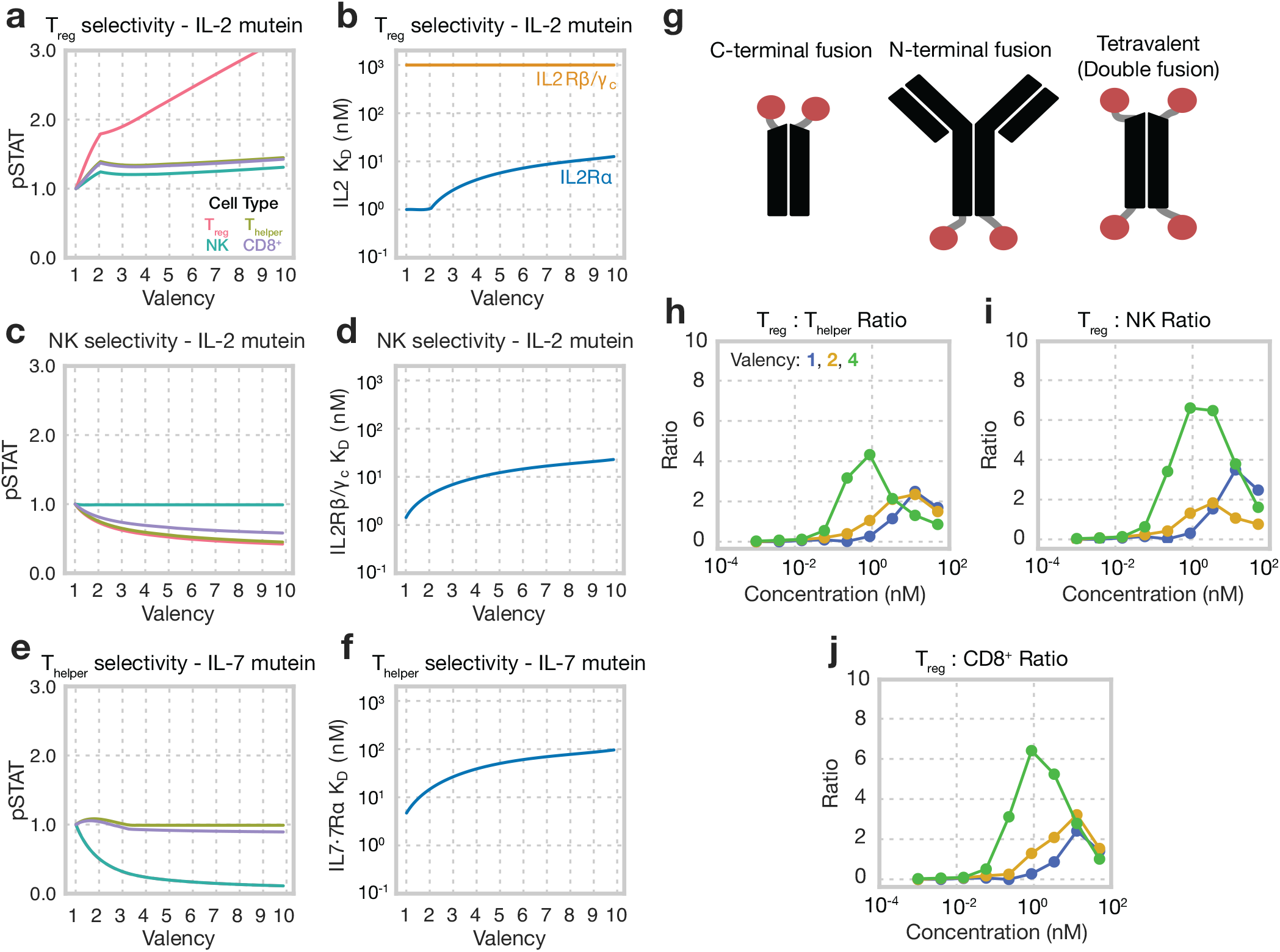
Multivalency enhances the selectivity of cytokine fusion proteins. **(a**,**c**,**e)** Signaling response of T_reg_, NK cells, and T_helper_ cells predicted for ligand of optimal selectivity at different valencies. Response predictions were normalized to each population’s response for the monovalent case. Selectivity for T_reg_s and NK cells were derived from IL-2 muteins, and selectivity for T_helper_s was calculated using IL-7 muteins. **(b**,**d**,**f)** Optimal receptor-ligand dissociation constants for each ligand optimized for selectivity. Mutein affinity for IL2Rα and IL2Rβ/γ_c_ was allowed to vary for IL-2 muteins, and affinity for IL-7Rα was allowed to vary for IL-7 muteins. Affinities were allowed to vary across K_D_s of 10 pM–1 µM while *K*^∗^ was fixed at its fitting optimum. All optimizations were performed using a concentration of 1 nM. Selectivity was calculated as the ratio of predicted pSTAT5 in target cells to the mean pSTAT5 predicted in off-target cells. **(g)** Schematic of multivalent IL-2 mutant design. **(h–j)** Ratio of STAT5 phosphorylation in T_reg_s to T_helper_s (h), NK cells (i), and CD8^+^ (j) cells at varying dosages for R38Q/H16N in various valency formats. Dots are representative of mean of experimental replicates (N=3).

We then explored whether IL-2 muteins lacking IL2Rα binding could selectively target NK cells, based on their uniquely high expression of IL2Rβ^15^, with similar results; IL-2 muteins of higher valency were predicted to be increasingly selective for activation of NK cells, so long as IL2Rβ/γ_c_ affinity was coordinately decreased (Fig. 5c,d). Finally, we explored whether multivalent IL-7 could be used to target T_helper_s, as they express high amounts of IL-7Rα (Fig. S1i). We again found that ligands of higher valency should achieve higher selectivity for these cells, but that the benefits of valency were less than the targeting of T_reg_s or NK cells using IL-2 mutants because CD8^+^ T cells have similar IL7Rα amounts (Fig. 5e). These benefits were again contingent on decreasing IL-7Rα affinity at higher valency (Fig. 5f).

To experimentally show that muteins of higher valency could be engineered to increase T_reg_ selectivity, we expressed and purified three Fc fusions of R38Q/H16N IL-2 in monovalent, bivalent, and tetravalent formats (Fig. S6). PBMCs from three donors were stimulated for 30 minutes and stained for cell type markers as well as pSTAT5. Tetravalent IL-2 was designed by Fc-fusing IL-2 muteins at both the C- and N-terminus and allowing the Fc to dimerize (Fig. 5g). R38Q/H16N was selected as the mutant closest to optimal binding affinities in tetravalent form, though further optimization is possible (Figs. 1b & 5b). As predicted, valency increased the responsiveness of both T_reg_s and off-target immune cells at each concentration (Fig. S7). However, the T_reg_ response increase far exceeded that in off-target cells; consequently, tetravalent R38Q/H16N was able to achieve much greater T_reg_ selectivity than bivalent and monovalent formats, two IL-2 fusions with already superior selectivity (Fig. 5h–j).

In total, these results show that valency beyond bivalency has unexplored potential for engineered cytokines with enhanced therapeutic potency alongside reduced toxicity. They also show the benefit of mechanistic modeling to guide ligand design, particularly when ligand affinity must be considered alongside other parameters such as valency.

### Bitargeted IL-2–Fc fusions demonstrate even greater T_reg_ selectivity

Through the CITE-seq data analysis, we found that IL2Rα was the optimal surface target for T_reg_ selectivity (Fig. 2i–m). This result was further strengthened when we integrated these data with our binding model and ligand optimization approach. We used the model to consider whether WT IL-2 fused to a selective binder for any surface markers could increase T_reg_ selectivity. WT IL-2 fusion to an IL2Rα binder was predicted to enhance T_reg_ selectivity over off-target immune cells (Fig. 6a/b). This was initially surprising because IL-2 itself binds IL2Rα but indicated to us that multivalent complexes provide the potential opportunity to decouple T_reg_ selectivity (by binding IL2Rα) from cytokine potency.

**Fig. 6.**
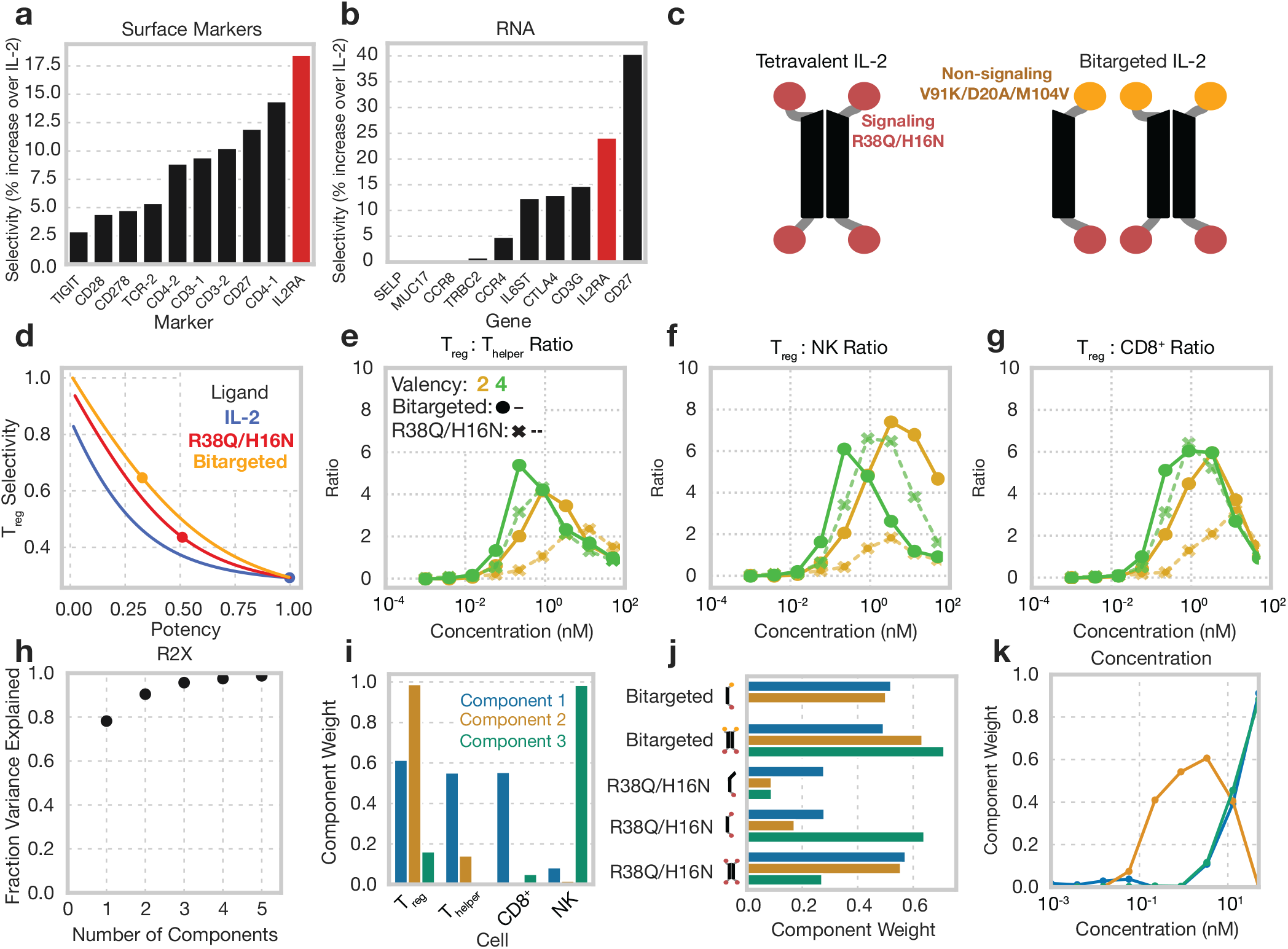
Asymmetric IL-2 mutants display even greater T_reg_ selectivity. **(a**,**b)** Predicted enhancements to T_reg_ selectivities for IL-2 mutants including a binding domain as calculated using CITE-seq surface marker data (a) or surface expressed RNA transcripts (b). Selectivity was calculated for T_reg_s against all other PBMC cells as the increase in average T_reg_ to off-target cell binding against WT IL-2 at a concentration of 0.1 nM. **(c)** Schematic of asymmetric IL-2 mutant design. **(d)** Predicted normalized T_reg_ selectivity displayed by bivalent WT IL-2, R38Q/H16N, and Bitargeted IL-2 across IL2Rβ affinities. **(e–g)** Ratio of STAT5 phosphorylation in T_reg_s to T_helper_s (e), NK cells (f), and CD8^+^ (g) cells at varying dosages for R38Q/H16N in various valency formats. Dots are representative of mean of experimental replicates (N=3). **(h–j)** Tensor factorization of signaling responses to R38Q/H16N and Bitargeted variants. **(h)** Percent variance reconstructed (R2X) versus the number of components used. **(i)** Component values representing cell type specificity. **(j)** Component values for each IL-2 mutant. **(k)** Component values representing the effect of concentration.

To decouple cell-selective binding from signaling response, we expressed an asymmetric Fc fusion including both a signaling-competent R38Q/H16N IL-2 and signaling-deficient V91K/D20A/M104V IL-2 with only IL2Rα binding (Fig. 6c, Fig. S6). V91K/D20A/M104V IL-2 is reported to effectively eliminate IL2Rβ binding while maintaining IL2Rα interaction^23^. We henceforth refer to this asymmetric construct as “bitargeted” IL-2 to reflect its inclusion of two IL-2 muteins with separate signaling and targeting roles. Bivalent bitargeted ligand was designed by introducing Fc mutations preventing Fc dimerization (Fig. 6c). Tetravalent bitargeted constructs were predicted to have greater T_reg_ specificity than their non-bitargeted counterparts for any IL2Rβ affinity, using either the IL2Rα affinity of WT or R38Q/H16N (Fig. 6d). We tested our bivalent and tetravalent bitargeted constructs by again stimulating PBMCs and quantifying their pSTAT5 responses. We found that both bivalent and tetravalent bitargeted ligand increased or maintained high potency in T_reg_ cells (Fig. S8). This potency translated to greater T_reg_ selectivity for both bivalent and tetravalent forms of bitargeted, both of which outperformed any previously characterized monovalent and bivalent IL-2 fusions, and modestly outperformed tetravalent R38Q/H16N (Fig. 6e–f).

We again applied non-negative CP decomposition of the R38Q/H16N and bitargeted ligand signaling responses to summarize our ligand engineering efforts (Fig. 6h–k). Three components captured >90% of the variation in the data. Here, T_reg_ responses were primarily represented by component 2, effector T cell responses by component 1, and NK cell responses by component 3 (Fig. 6i). Valency again determined ligand component 2 weight most potently, with both tetravalent constructs, bitargeted or R38Q/H16N, demonstrating highest component 2 weight. However, while the bivalent bitargeted construct demonstrated no NK cell activity (component 3), both tetravalent constructs had some NK activity (Fig. 6j). Thus, our experimental results demonstrate that tradeoffs still exist between T_reg_ potency and selectivity. As before, T_reg_ selectivity was maximized at low and intermediate dosages (Fig. 6k). Despite these tradeoffs, the selectivity demonstrated by both model-guided “cis”-targeting or higher valency fusion formats vastly improved upon the selectivity possible with existing approaches.

## DISCUSSION

Here, we systematically explored how ligand properties determine signaling response and specificity across 13 engineered IL-2 variants (Fig. 1, 2). Our study is unique in its inclusion of clinically relevant muteins alongside variation in Fc fusion format. Dimensionality reduction in tensor form identified how ligand properties modulate response, revealing that multivalent cytokines have unique specificity advantages (Fig. 3). Using a multivalent binding model, we uncovered that this unique specificity arises from surface binding avidity effects (Fig. 4). This model indicated that cytokines of higher valency can offer even greater cell type selectivity (Fig. 5), and we demonstrated this strategy experimentally by expressing tetravalent IL-2 fusions with greater T_reg_ selectivity than current state-of-the-art monovalent or bivalent affinity muteins. Finally, we uncovered that IL2Rα itself is the optimal target for designing T_reg_-selective binding, and that cis-targeting can be designed into multivalent IL-2 fusions through asymmetric tetravalent IL-2 fusions, again improving on signaling selectivity (Fig. 6). In total, our results show that not only do multivalency and cis targeting of IL2Rα improve T_reg_ selectivity, but that these paired strategies represent the only known method for overcoming the selectivity-potency tradeoff faced by T_reg_-selective muteins^18^.

Our results have clear implications for the design of T_reg_-directed IL-2 therapies, an area of enormous interest for the treatment and management of autoimmune diseases^22,38^. We showed computationally and experimentally that multivalency and bitargeting can enhance IL-2 T_reg_ selectivity for potential use in clinical settings, where IL-2 based therapies have traditionally struggled^39^. Engineering valency requires precise compensatory adjustments in the ligand affinity (Fig. 5); given that we limited our experiments to pre-existing muteins, we expect that our selectivity gains might be improved even further by identifying muteins with optimal affinities. Our multivalent and bitargeted designs will additionally need to be tested *in vivo* to see that these selectivity gains translate to the *in vivo* setting^23^. T_reg_ selectivity is central to the mechanism of action for these therapies, and so we expect that these benefits to selectivity will improve therapeutic properties in several ways: more potent activation of signaling in T_reg_s without off-target effects may improve the potency of these therapies and the breadth of applications^18,40,41^; reduced toxicity may allow for more routine use with minimal patient monitoring^14^. The superior selectivity offered by engineered multivalent ligands will likely further increase their *in vivo* pharmacokinetic lifetimes, in turn requiring less frequent dosing, as most drug clearance occurs via receptor-mediated endocytosis in off-target populations^21,42–44^. However, known differences exist in IL-2 receptor expression between humans and mice, so we do not expect that results from murine models would be a reliable indication for the comparative advantages of these molecules^15,16^, and primate studies are outside the scope of the present work.

Heterospecificity, in our case exploited through bitargeting, opens a whole range of new possibilities through its ability to decouple the targeting and signaling properties of cytokine therapy and/or combine synergistic signals. This capability has been demonstrated through bispecific antibodies previously, and through the design of cis-targeted cytokine-antibody fusions. However, we showed that, unlike other immune cells, T_reg_s do not express any surface marker more selective than IL2Rα (Fig. 2i-l). Consequently, there are no alternative targeting options to our approach of using the IL-2 receptors themselves (Fig. 6). Beyond our results, heterospecificity creates opportunities for synergistic receptor agonism. For example, PD-1 cis-targeting with IL-2 increases the stemness of CD8^+^ T cells and consequently their tumor killing capacity^45,46^. While IL-2 has been employed as a therapy because of its T_reg_ selectivity, there is no reason to believe that the cytokine’s signaling effects are optimal for enhancing T_reg_ suppressive activities. In fact, with cell therapies, where selectivity is not a concern and non-natural cytokine receptors can be introduced, other cytokine signaling such as IL-9 is qualitatively more effective than IL-2 at promoting cytotoxic T cell function^47,48^. Thus, one possibility enabled by bitargeting is potentially plug-and-play combinations of one or more cytokines that are more capable than IL-2 of driving desirable T_reg_ properties, made T_reg_-selective through their fusion to multivalent IL2Rα-targeting complexes^49–51^. More systems-level research into the signaling regulation of T_reg_ proliferation and suppressive activities, and comparisons to other cytokines beyond IL-2, is needed to develop these possibilities.

Most generally, our results demonstrate the value of computationally directed biologics design, particularly for fusion constructs incorporating more than one binding moiety. The design of these modular ligands leads to a combinatorial explosion of ligand configurations^52^. Furthermore, multivalent ligands have several documented effects, including altered signal transduction^53,54^, binding avidity, pharmacokinetics^55^, and intracellular trafficking^56^. While valency has been extensively applied as a means to introduce binding selectivity based on receptor density^57,58^, multiple receptors, ligands subunits with varied targeting, and differences in signaling effects lead to additional complexity. For instance, while we found that the contribution of multivalency was explained primarily through avidity effects (Fig. 4), bitargeting as a strategy arises through differences in the signaling capacity of IL2Rα versus IL2Rβ. The approach here— computationally designed, multivalent, bitargeted ligands for enhanced therapeutic selectivity—has widespread application to other receptor-ligand pathways, including IL-4/IL-13, bone morphogenic proteins, and the TNF cytokines^52,59,60^. There are likely still other design strategies to be found across these many signaling pathway structures.

## MATERIALS AND METHODS

### EXPERIMENTAL METHODS

Receptor abundance quantitation, octet binding assays, expression of recombinant bivalent and monovalent IL-2 muteins (Fig. 1–4), and measurement of those muteins’ signaling in PBMCs were performed as described in Farhat *et al*.^15^.

### Receptor abundance quantitation

Receptor quantitation data was gathered as described previously in Farhat et al.^15^; the preprocessing of fluorescence measurements, population gating, and receptor abundance calculations were performed using these data. To quantify the number of antibodies bound to cells and to standard beads, the fluorescence intensity of isotype controls was subtracted from the signal from matched receptor stains and then calibrated using the two lowest quantitation standards. Cell gating was conducted as shown in Fig. S1. The geometric means of replicates were calculated to summarize the results.

### pSTAT5 Measurement of IL-2 and -15 Signaling in PBMCs

Cryopreserved PBMCs (UCLA Virology Core) were thawed to room temperature and slowly diluted with 9 mL pre-warmed RPMI-1640 (Corning, 10040CV) supplemented with 10% FBS (VWR, 97068-091, lot#029K20) and Penicillin/Streptomycin (Gibco, 15140122). Media was removed, and cells were brought to 3×10^6^ cells/mL, distributed at 300,000 cells per well in a 96-well V-bottom plate, and allowed to recover 2 hrs at 37℃ in an incubator at 5% CO_2_. IL-2 (R&D Systems, 202-IL-010) or IL-15 (R&D Systems, 247-ILB-025) were diluted in RPMI-1640 in the absence of FBS. These dilutions were then added to the concentrations indicated. To quantify STAT5 phosphorylation, the media was taken away, and cells were fixed using 100 µL of 10% formalin (Fisher Scientific, SF100-4) for 15 mins at room temperature. Formalin was removed from the cells, and the PBMCs were placed on ice.

They were then suspended in 50 µL of cold methanol (−30℃). PBMCs were then kept at - 30℃ overnight. PBSA was used to wash the cells twice. The cells were then split into two identical plates and stained with fluorescent antibodies for 1 hr at room temperature in darkness using 50 µL of antibody panels 4 and 5 per well. Cells were suspended in 100 µL PBSA per well, and beads to 50 µL, and analyzed on an IntelliCyt iQue Screener PLUS with VBR configuration (Sartorius) using a sip time of 35 secs and beads 30 secs. Compensation of measured fluorescent values was calculated as detailed above. Gating of cell populations was performed as shown in Fig. S1, and the median pSTAT5 level was calculated for each population in each well.

### Recombinant proteins

The Expi293 expression system was used to express IL-2/Fc fusion proteins. Expression was conducted as prescribed by the manufacturer instructions (Thermo Scientific). Proteins were formulated as the Fc of human IgG1 fused at its N- or C-terminus to human IL-2 using a (G_4_S)_4_ linker. C-terminal lysine residues of human IgG1 were not included in C-terminal fusions. The AviTag sequence GLNDIFEAQKIEWHE was added to the Fc terminus which did not contain IL-2. Fc mutations which prevented dimerization were introduced into the Fc sequence for monovalent muteins^61^. MabSelect resin (GE Healthcare) was used to purify protein. Biotinylation of proteins was conducted using BirA enzyme (BPS Biosciences) according to manufacturer instructions. Extensive buffer-exchanging into phosphate buffered saline (PBS) was conducted using Amicon 10 kDa spin concentrators (EMD Millipore). The sequence which was used to express the IL2Rβ/γ Fc heterodimer was the same as that of a reported, active heterodimeric molecule (patent application US20150218260A1); a (G4S)2 linker was added between the Fc portion and each receptor ectodomain. The Expi293 system was used to express the protein, which was subsequently purified on MabSelect resin as above. The IL2Rα ectodomain was generated to include a C-terminal 6xHis tag and then purified on Nickel-NTA spin columns (Qiagen) according to manufacturer instructions.

### pSTAT5 Measurement of Tetravalent IL-2 Signaling in PBMCs

Cryopreserved PBMCs (UCLA Virology Core) were thawed to room temperature and slowly diluted with 9 mL pre-warmed RPMI-1640 (Corning, 10040CV) supplemented with 10% FBS (VWR, 97068-091, lot#029K20) and Penicillin/Streptomycin (Gibco, 15140122). Media was removed, and cells were brought to 3×10^6^ cells/mL, distributed at 300,000 cells per well in a 96-well V-bottom plate, and allowed to recover 2 hrs at 37℃ in an incubator at 5% CO_2_. IL-2 (Peprotech, 200-02-50μg) and tetravalent IL-2 (expressed and purified as described below) were diluted in RPMI-1640 without FBS and added to the indicated concentrations. To measure pSTAT5, media was removed, and cells fixed in 100 µL of 4% paraformaldehyde (PFA, Election Microscopy Sciences, 15714) diluted in PBS for 15 mins at room temperature.

PFA was removed, cells were gently suspended in 100 µL of cold methanol (−30℃). Cells were stored overnight at -30℃, and then washed twice with 0.1% bovine serum albumin (BSA, Sigma-Aldrich, B4287-25G) in PBS (PBSA), and stained 1 hr at room temperature in darkness using antibody panel X with 40 µL per well. Cells were then washed twice with 0.1% PBSA and resuspended in 150 µL PBSA per well. Cells were analyzed on a BD FACSCelesta flow cytometer. Populations were gated as shown in **Fig. S1**, and the median pSTAT5 level was extracted for each population in each well.

### Tetravalent IL-2 Expression

Proteins were expressed as human IgG1 Fc-fused at the N- or C-terminus to mutant human IL-2 through a flexible (G4S)4 linker. C-terminal fusions omitted the C-terminal lysine residue of human IgG1. In bivalent variants, Fc mutations to prevent dimerization were introduced into the Fc sequence. Each IL-2 fused via the 20 amino acid long linker to the Fc domain contained R38Q and H16N mutations to reduce the IL-2’s affinity with which it binds IL2Rβ. Plasmid DNA prepared by maxi-prep (Qiagen, 12162) were transfected into adherent HEK293T cells using Lipofectamine 3000 (Thermo-Fisher, L3000008) in 15 cm dishes in DMEM (Corning, 15017CV) supplemented with GlutaMax (Gibco, 35050061) and 10% FBS. Media was exchanged after 24 hrs with fresh DMEM supplemented with GlutaMax and 5% ultra-low IgG FBS (Thermo-Fisher, A3381901). Media was harvested after an additional 72 hrs. Media was incubated in the presence of Protein A/G Plus Agarose resin (Santa Cruz Biotechnology, sc-2003) overnight. The following day, the media-resin mixture was centrifuged, and the supernatant discarded. Resin was washed with PBS five times or until protein was no longer detected in supernatant by UV-Vis using a NanoDrop One Spectrophotometer (Thermo-Fisher, ND-ONE-W). IL-2 was eluted from resin using 0.1M glycine, pH 2.3, into 2M Tris-HCl, pH 8. IL-2 was then buffer exchanged into PBS for storage at -80℃. Concentration was determined by BCA assay and confirmed using an IgG1 ELISA.

### Octet binding assays

An Octet RED384 (ForteBio) was used to measure the binding affinity of each IL-2 mutein. Monomeric, biotintylated IL-2/Fc fusion proteins were loaded to Streptavidin biosensors (ForteBio) at roughly 10% of saturation point and allowed to equilibrate for 10 min in PBS + 0.1% bovine serum albumin (BSA). Up to 40 min of association time in IL2Rβ/γ titrated in 2x steps from 400 nM to 6.25 nM, or IL2Rα from 25 nM to 20 pM, which was followed by dissociation in PBS + 0.1% BSA. A zero-concentration was included in each measurement and served as a negative control/reference signal. The affinity quantification experiments were performed in quadruplicate across two days. Binding of IL-2 to IL2Rα on its own did not fit to a simple binding model; K_D_ was calculated using equilibrium binding within each assay for this case. IL2Rβ/γ binding data fit a 1:1 binding model; thus, in these cases on-rate (k_on_), off-rate (k_off_) and K_D_ were determined by fitting to the entire binding curve. The average of each kinetic parameter across all concentrations with detectable binding (typically 12.5 nM and above) was used to calculate K_D_.

### Statistical analysis

The number of replicates performed for each experimental measurement, and the values of confidence intervals are described in corresponding figure captions. N is used to describe the number of times a particular experiment was performed. Flow cytometry experiments performed using initial panel of monovalent and bivalent cytokines (Figs. 1–4) were performed on hPBMCs were conducted using separate experimental replicates on cells gathered from a single donor. Each replicate of the flow cytometry signaling experiments in figures 5 and 6 were conducted using hPBMCs from different donors. To quantify population-level flow cytometry measurements for both signaling and receptor quantitation experiments, the mean fluorescent intensity (MFI) of a gated population was measured.

Compensation to remove fluorescent spectral overlap was performed for each experimental measurement. Subtraction of either negative controls or cells treated with isotype antibodies was performed on signaling and receptor quantitation data respectively to remove background signal. Cells which were measured to display fluorescent intensities above 1,000,000 were excluded from analysis during signaling experiments. Pearson correlation coefficients (R^2^) values were used to describe model accuracy when predicting signaling response to IL-2 and IL-2 muteins. The K_x*_ parameter was fit with least-squares fitting using the Broyden–Fletcher–Goldfarb–Shanno minimization algorithm as implemented in SciPy.

## MODELING

### Binding model

The model was formulated as described in Tan *et al*^32^. The monomer composition of a ligand complex was represented by a vector 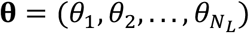, where each *θ*_*i*_ was the fraction of monomer ligand type *i* out of all monomers on that complex. Let *C*_**θ**_ be the proportion of the **θ** complexes in all ligand complexes, and *Θ* be the set of all possible **θ**’s. We have ∑_**θ**∈Θ_ *C*_**θ**_ = 1.

The binding between a ligand complex and a cell expressing several types of receptors can be represented by a series of *q*_*ij*_. The relationship between *q*_*ij*_’s and *θ*_*i*_ is given by 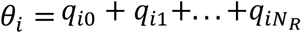. Let the vector 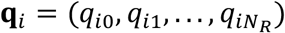, and the corresponding **θ** of a binding configuration **q** be **θ**(**q**). For all *i* in {1,2, . . ., *N*_*L*_}, we define 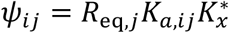 where *j* = {1,2, . . ., *N*_*R*_} and *Ψ*_*i*0_ = 1. The relative number of complexes bound to a cell with configuration **q** at equilibrium is

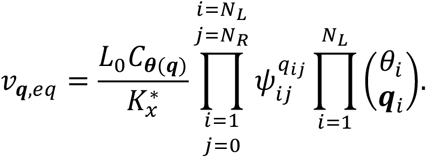

Then we can calculate the relative amount of bound receptor *n* as

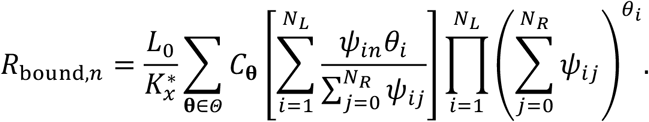

By mass conservation, (*R*_tot,n_ = *R*_eq,n_ + *R*_bound,n_), we can solve *R*_eq,n_ numerically for each type of receptor.

### Application of multivalent binding model to IL-2 signaling pathway

Each IL-2 molecule was allowed to bind to one free IL2Rα and one IL2Rβ/γ_c_ receptor. Initial IL-2-receptor association proceeds with the known kinetics of monomeric ligand-receptor interaction (Table S1). Subsequent ligand-receptor binding interactions then proceed with an association constant proportional to available receptor abundance and affinity multiplied by the scaling constant, 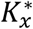, as described above. To predict pSTAT5 response to IL-2 stimulation, we assumed that pSTAT5 is proportional to the amount of IL-2-bound IL2Rβ/γ_c_, as complexes which contain these species actively signal through the JAK/STAT pathway. Scaling factors converting from predicted active signaling species to pSTAT5 abundance were fit to experimental data on a per-experiment and cell type basis. A single 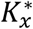 valuewas fit for all experiments and cell types.

### CITE-seq marker selectivity analysis

To assist in identifying possible methods through which to increase IL-2 selectivity towards T_reg_s, a publicly available Cellular Indexing of Transcriptomes and Epitopes by sequencing (CITE-seq) dataset containing data gathered from human PBMCs was analyzed^28^. Only RNA transcripts encoding cell membrane extracellular-facing proteins were included. We first analyzed the data by determining the Wasserstein distance and Kullback-Leibler divergence of markers and RNA measured in T_reg_s against the distribution of these markers displayed by all other cells. We also analyzed the data using a ridge classification model, where all markers and RNA sequences were used by the model to distinguish between T_reg_s and all other cell types.

Markers of interest were then used in conjunction with the binding model to determine whether they could confer selectivity, using the CITE-seq data to inform the number of markers per cell. Conversion factors for calculating marker abundance from CITE-seq marker and mRNA reads were estimated using proportional conversions from the data to previously experimentally determined marker abundances^15^. Single cell marker abundances were calculated for 1000 cells at a time, and the ratio of T_reg_ signaling to off-target signaling was calculated. To simulate bispecific binding, two distinct binding domains for each ligand were modeled, one for IL2, with affinity for IL2Ra and IL2RB, and the other for the marker of interest. An optimization function was used to calculate the binding affinity which would yield the best selectivity per marker. After finding IL2Rα to be the optimal epitope for increasing selectivity, we sought to explore the effects of increasing valency by doubling the number of binding domains per ligand.

### Tensor Factorization

Before decomposition, the signaling response data was background subtracted and variance scaled across each cell population. Non-negative canonical polyadic decomposition was performed using the Python package TensorLy, using the HALS algorithm with non-negative SVD initialization^62^.

## Acknowledgements

The authors thank Cori Posner for contributing to initial profiling studies that enabled the study.

## Funding

This work was partly supported by startup funds from UCLA Engineering and by NIH U01-AI148119 to A.S.M.

## Author contributions statement

A.S.M. conceived of the study. S.D.T. performed the PBMC experiments with the IL-2 fusion proteins. A.S.M, B.O.J., E.M.S., and P.C.E. performed the computational analysis. All authors helped to design experiments and/or analyze the data. All authors contributed to writing the paper.

## Competing interests

A.S.M. has filed a patent PCT/US22/35711 on the use of multivalent cytokines to enhance cell type-selective responses. A.S.M., B.O.J., and P.C.E. have filed a provisional patent on the use of bitargeting for engineered cytokine responses.

## Data Sharing Plan

All analysis was implemented in Python v3.10 and can be found at https://github.com/meyer-lab/gc-valent, release 1.0, along with all the experimental data.

## Notes

https://github.com/meyer-lab/gc-valent

### Summary of Updates

We have greatly expanded the paper with experimental validation of tetravalent Fc fusions, and a new bitargeting approach. All the figures have been revised.

