## Supplemental Materials for "Multivalent, asymmetric IL-2-Fc fusions provide optimally enhanced regulatory T cell selectivity"

### Supplementary Figures

**Table S1. IL-2 variants' affinities for IL-2R subunits.**

| Ligand | IL-2Rα $K_D$ (nM) | IL-2Rβ/$\gamma_c$ $K_D$ (nM) |
|---|---|---|
| WT IL-2 | 10.0 | 0.133 |
| WT N-term | 0.19 | 5.30 |
| WT C-term | 0.54 | 3.04 |
| V91K C-term | 0.69 | 7.56 |
| R38Q N-term | 0.71 | 4.00 |
| F42Q N-Term | 9.48 | 2.81 |
| N88D C-term | 1.01 | 24.0 |
| H16N N-term | 0.43 | 22.4 |
| R38Q/H16N | 0.71 | 22.4 |

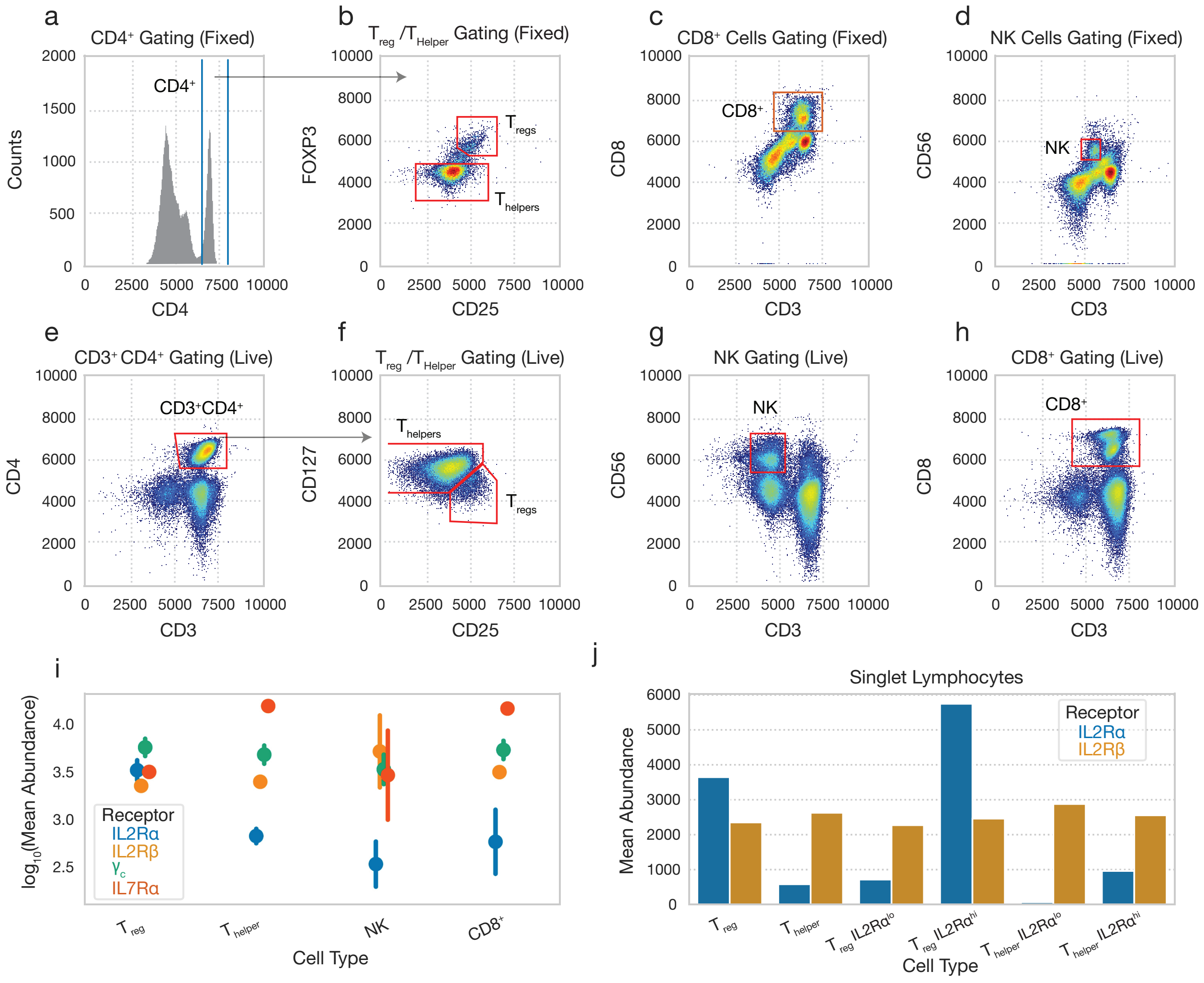


**Fig. S1. Receptor quantification and gating of PBMC-derived immune cell types.** **(A&B)** Gating for fixed $T_{helper}$ and $T_{reg}$ cells during pSTAT5 quantification. **(C&D)** Fixed CD8+ T cell and NK cell gating. **(E&F)** Gating for live $T_{helper}$ and $T_{reg}$ cells during receptor quantification. **(G)** Live cell NK cell gating. **(H)** Live cell CD8+ cell gating. **(I)** Receptor quantification for each cell type. Experiments were performed in quadruplicate (N=4). **(J)** IL2Rα and IL2Rβ abundances on IL-2Rα high and low $T_{reg}$ and $T_{helper}$ populations. Cells were binned using three evenly logarithmically spaced separations between $5^{th}$ and $95^{th}$ percentile of IL-2Rα abundance.

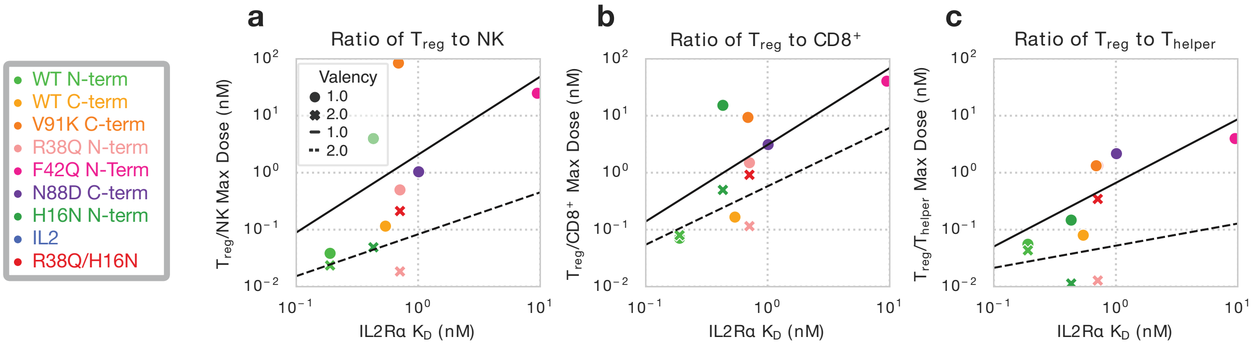


**Fig. S2. Concentration of optimum ligand selectivity is a function of IL2Rα affinity. (a-c)** Location of the concentration at which the ratio of $T_{reg}$ to NK (a), CD8$^+$ (b), and $T_{helper}$ (c) activity is maximized vs. IL2Rα affinity. Lines were fit to monovalent (solid) and bivalent (dotted).

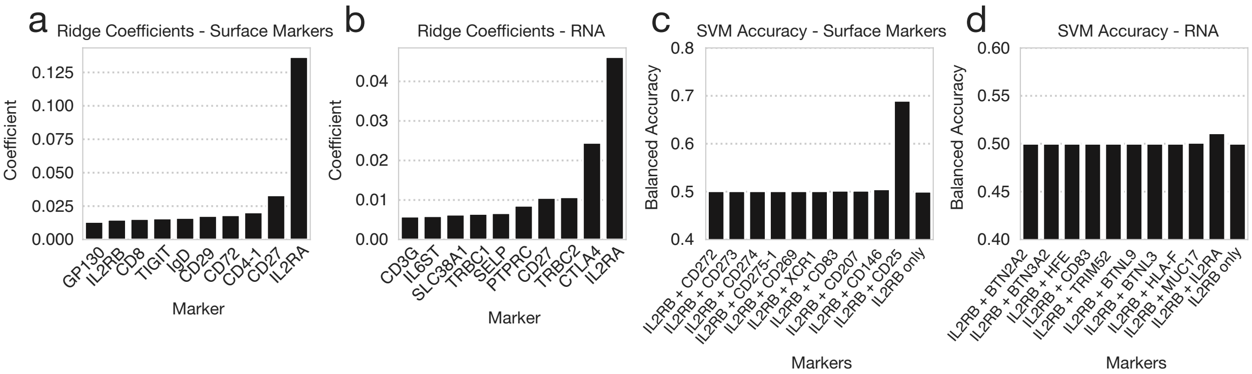


**Fig. S3. Linear and non-linear classification algorithms identify IL2Rα as most unique marker on $T_{reg}$s. (a,b)** Largest marker coefficients determined by fitting a RIDGE-classifier to CITE-seq surface marker data (a) and mRNA data (b). Model was fit to identify $T_{reg}$s using a one-vs.-all approach. **(c,d)** Largest $T_{reg}$ identification accuracies of Support Vector Classifier fit using IL2Rβ and one other marker using surface marker (c) and RNA (d) data. Accuracy is reported as Balanced Accuracy.

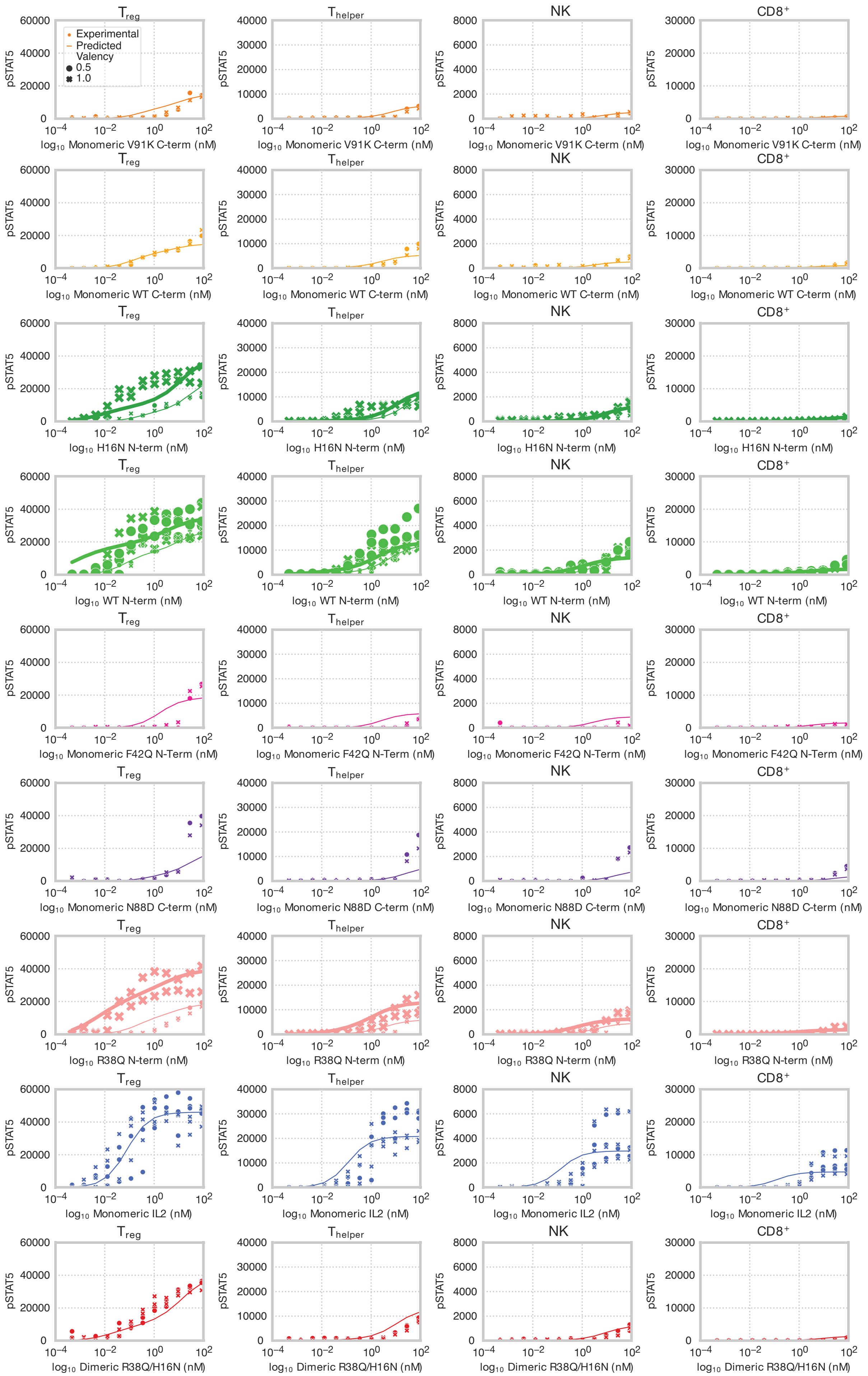

Experimental
Predicted
Valency
0.5
1.0
Treg
Thelper
NK
CD8+
pSTAT5
log10 Monomeric V91K C-term (nM)
log10 Monomeric WT C-term (nM)
log10 H16N N-term (nM)
log10 WT N-term (nM)
log10 Monomeric F42Q N-Term (nM)
log10 Monomeric N88D C-term (nM)
log10 R38Q N-term (nM)
log10 Monomeric IL2 (nM)
log10 Dimeric R38Q/H16N (nM)

**Fig. S4. Full panel of predicted versus experimental immune cell type responses to monomeric and dimeric IL-2 muteins.** Dots represent flow cytometry measurements and lines represent pSTAT response predicted by model. Experimental pSTAT measurements are shown for 0.5- and 1-hour timepoints. Predictions and experiments are shown for $T_{reg}$s, $T_{helper}$s, NK and CD8 cells. Each point is representative of one experimental output (N=1). Shaded regions are indicative of standard error prediction when scalar factor converting between signaling complexes and MFI was fit to multiple experiments.

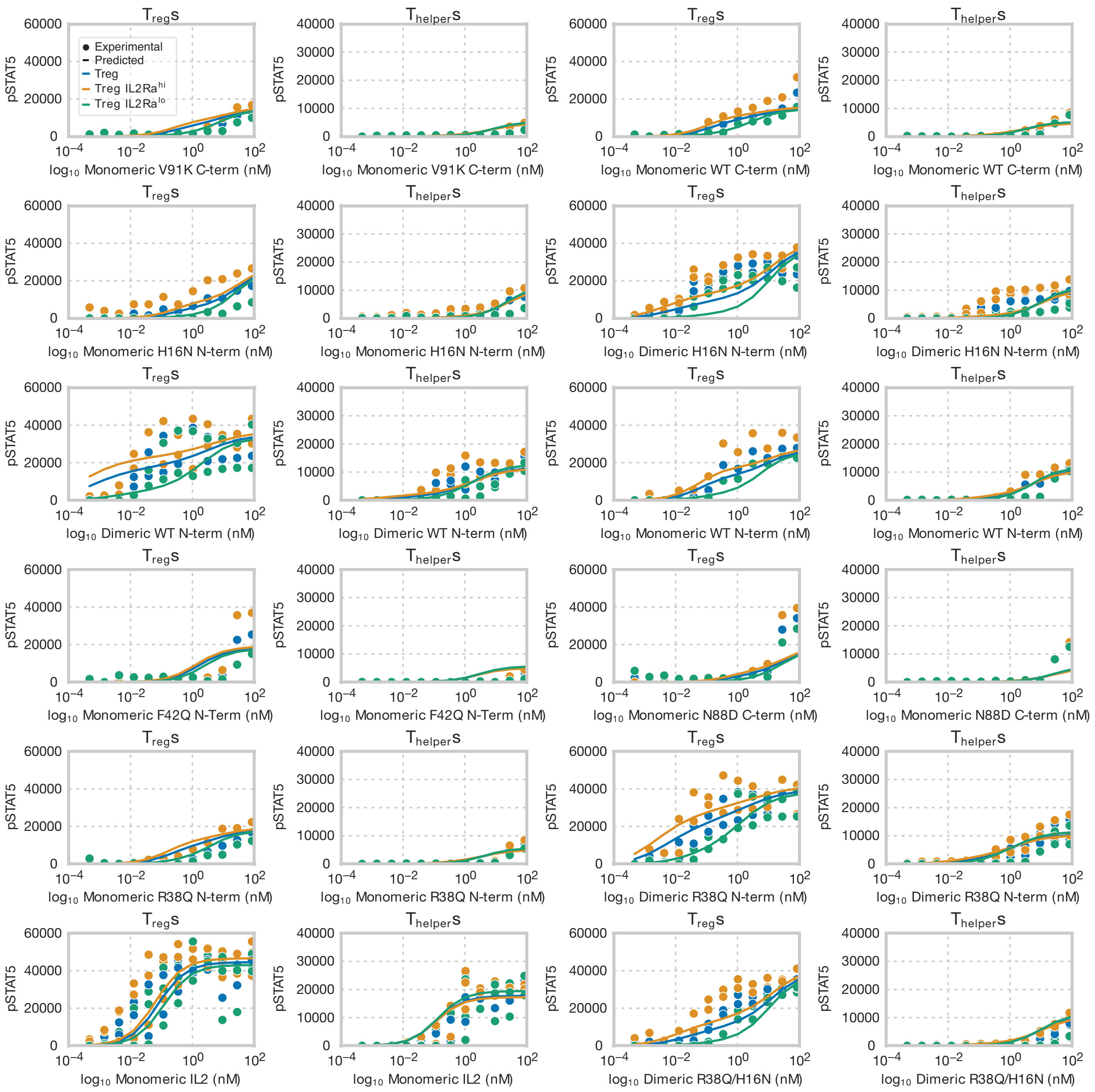


**Fig. S5. Full panel of predicted versus experimental IL-2Rα high, medium, and low $T_{reg}$ and $T_{helper}$ responses to monomeric and dimeric IL-2 muteins.** Dots represent flow cytometry measurements and lines represent pSTAT response predicted by model. Experimental pSTAT measurements are shown for 0.5- and 1-hour timepoints. Predictions and experiments are shown for $T_{reg}$s, $T_{helper}$s binned by their IL-2Rα abundances. Each point is representative of one experimental output (N=1). Shaded regions are indicative of standard error prediction when scalar factor converting between signaling complexes and MFI was fit to multiple experiments.

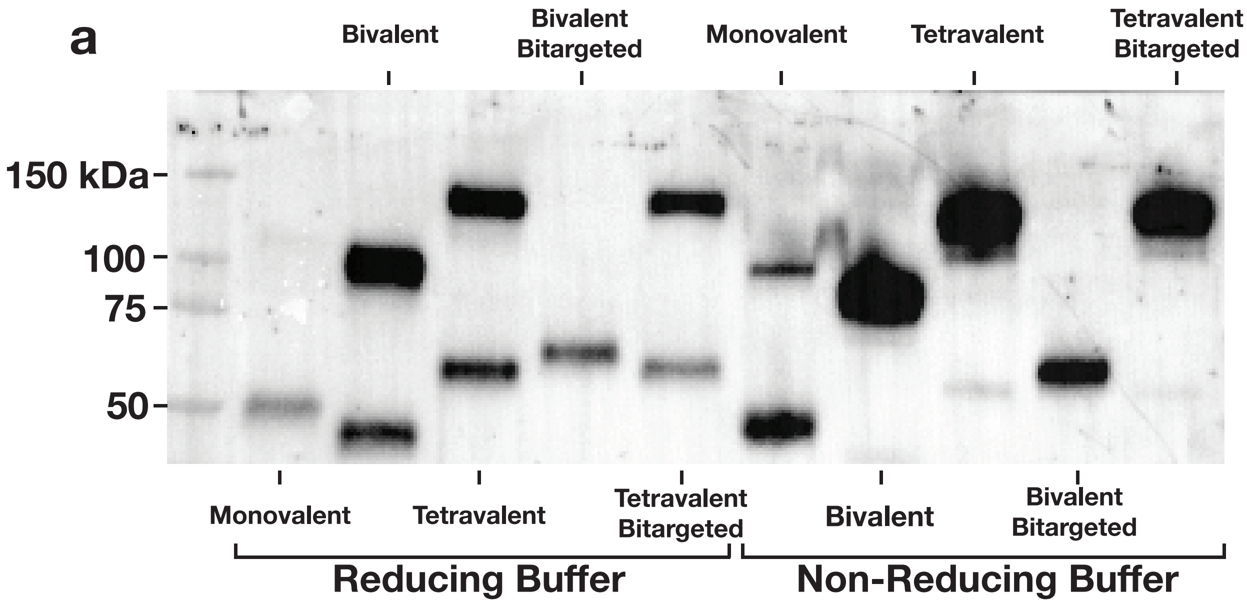


**Fig. S6. Western blot of multivalent IL-2 constructs.** Western blot of monovalent, bivalent, tetravalent R38Q/H16N (lanes 1–3 and 6–8) and bivalent and tetravalent Bitargeted IL-2 (lanes 4–5 and 9–10). Samples in lanes 1–5 were run in reducing buffer, and lanes 6–10 were run in non-reducing buffer. Blot was stained with a primary anti-human-IL-2 antibody.

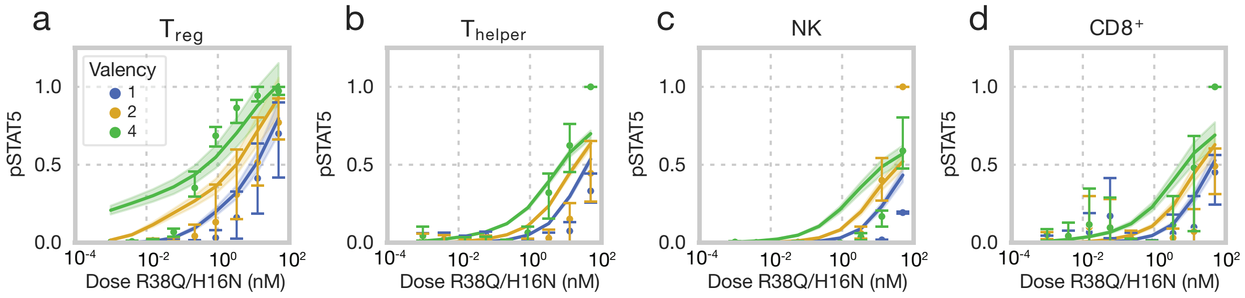


**Fig. S7. Full panel of predicted and experimental responses to R38Q/H16N multivalent mutants.** Responses of human $T_{reg}$s (a), $T_{helper}$s (b), NK (c), and $CD8^+$ (d) cells, measured by STAT5 phosphorylation, in response to varying dosages of R38Q/H16N in various valency formats. Cells were stimulated with cytokine for 30 minutes and signaling was normalized to the largest signaling response for each cell type and donor. Points are representative of experimental results (N=3), error bars represent experimental standard deviation, and lines represent model-predicted responses.

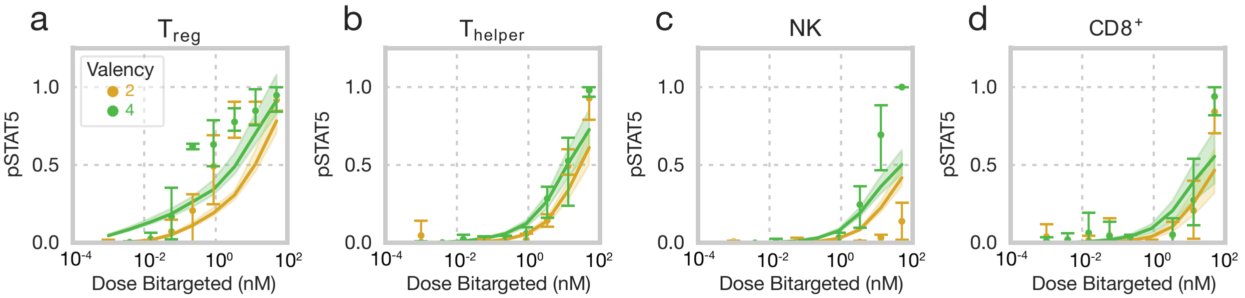


**Fig. S8. Full panel of predicted and experimental responses to bitargeted multivalent mutants.** Responses of human $T_{reg}$s (a), $T_{helper}$s (b), NK (c), and $CD8^+$ (d) cells, measured by STAT5 phosphorylation, in response to varying dosages of Live/Dead IL-2 in various valency formats. Cells were stimulated with cytokine for 30 minutes and signaling was normalized to the largest signaling response for each cell type and donor. Points are representative of experimental results (N=3), error bars represent experimental standard deviation, and lines represent model-predicted responses.
